# Transfer Learning from Nucleus Detection to Classification in Histopathology Images

**DOI:** 10.1101/530113

**Authors:** Safoora Yousefi, Yao Nie

## Abstract

Despite significant recent success, modern computer vision techniques such as Convolutional Neural Networks (CNNs) are expensive to apply to cell-level prediction problems in histopathology images due to difficulties in providing cell-level supervision. This work explores the transferability of features learned by an object detection CNN (Faster R-CNN) to nucleus classification in histopathology images. We detect nuclei in these images using class-agnostic models trained on small annotated patches, and use the CNN representations of detected nuclei to cluster and classify them. We show that with a small training dataset, the proposed pipeline can achieve superior nucleus detection and classification performance, and generalizes well to unseen stain types.

## 1 INTRODUCTION

Nucleus detection is a core task in computational pathology and refers to the identification and localization of individual cells in histopathological images. Such images come in varied sizes, resolutions, stain types, and are often crowded with visually heterogeneous, overlapping cells. Among the challenges that are commonly faced in nucleus detection from these images are the difficulty of obtaining ground truth annotations, and transferring a model tuned for a specific resolution or stain to a dataset with different characteristics.

We approach nucleus detection as the general object detection problem where the goal is to find and classify a variable number of objects on an image. Object detection is a well studied problem in machine learning and computer vision where recently deep convolutional neural networks have shattered performance benchmarks. However, most of the advances made in this field assume availability of large training datasets, and are designed to handle relatively small images with a few objects in them, so applying these algorithms to whole slide images is not straightforward.

Building upon the Faster-RCNN deep object detection model [1] we propose an efficient nucleus classification pipeline that can be used for AI-assisted ground truth annotation. We train a Faster-RCNN for class-agnostic nucleus detection, and use the learned model for detection of and feature extraction from new classes of nuclei. Extracted features are later used to cluster nuclei into homogeneous subsets that can be labeled in bulk by human annotator, reducing the burden of annotation from an order of number of nuclei to the number of clusters. We demonstrate the ability of the learned model to detect nuclei in datasets with mixed sizes and resolutions, and to generalize to stain types that it has not seen during training.

## 2 RELATED WORK

Adaptive nucleus localization and classification has been approached previously as object detection, semantic segmentation, or regression. Regression random forests were used in [2] to predict nucleus centers. Object counting regression methods have also been applied to nucleus detection [3]. CNNs were used for supervised nucleus segmentation in microscopy images [4, 5] and supervised nucleus detection and classification in histopathology images [6].

CNNs are known to learn features that are generalizable to new tasks, and image representations derived from pretrained CNNs have become the state of the art in computer vision tasks such as instance retrieval [7] and cell classification [8]. To our knowledge, this is the first attempt to measure the transferability of nucleus detection CNN features to nucleus classification task.

## 3 DATASETS

**Pre-training dataset**: We used a Faster-RCNN model released along with the Tensorflow Object Detection API [9]. The model was pre-trained on the COCO dataset [10] which is a large scale general object detection dataset with 300K images from 80 categories of animals, furniture, vehicles, etc.

**Fine-tuning datasets:** In order to familiarize the pre-trained Faster-RCNN with the unique characteristics of histopathology images, we fine-tuned it on the following immunohistochemistry (IHC) bright field image datasets annotated by our trained research associates with class-agnostic bounding box annotations. These images are originally several thousands of pixels in each dimension, therefore, to facilitate annotation we extracted random patches of 64×64 or 128×128 pixels from them (see Table 1).

**Table 1.**
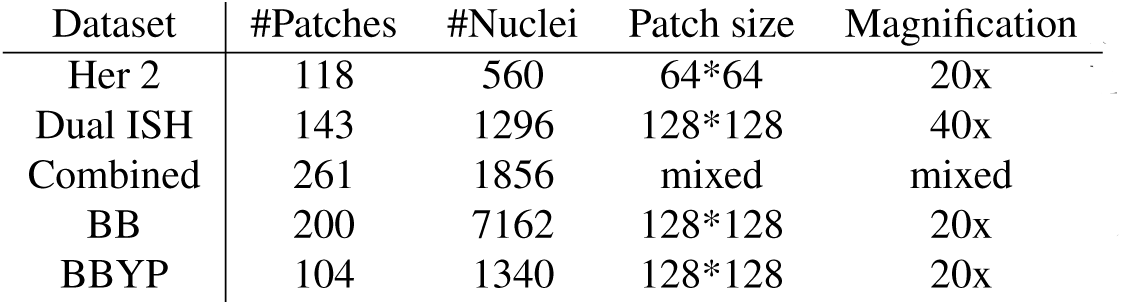
Summary of training data used for fine-tuning of Faster-RCNN.

| Dataset | #Patches | #Nuclei | Patch size | Magnification |
| --- | --- | --- | --- | --- |
| Her 2 | 118 | 560 | 64*64 | 20x |
| Dual ISH | 143 | 1296 | 128*128 | 40x |
| Combined | 261 | 1856 | mixed | mixed |
| BB | 200 | 7162 | 128*128 | 20x |
| BBYP | 104 | 1340 | 128*128 | 20x |

- Her2: Human epidermal growth factor receptor 2 membrane stained cells.
- Dual ISH: Nuclei with dual in situ hybridization.
- Combined: Combination of Her2 and Dual ISH datasets.

**Classification datasets**: Two datasets with both bounding box annotations and class labels were used for classification performance evaluation in transfer learning experiments:

- BB: 7162 two-class annotations of Hematoxylin and DAB stained nuclei.
- BBYP: IHC Duplex stained images containing 1340 annotations that fall into 5 imbalanced classes with number of members ranging from 10 to 1828.

## 4 METHODS

We train several models for different experiments. Each experiment consists of the following steps: fine-tuning a pretrained Faster RCNN model on one of our fine-tuning datasets for class-agnostic nucleus detection, feeding a new dataset to the fine-tuned model for inference, extracting detected nuclei representations from various layers, clustering the detected nuclei based on their extracted representations, and measuring clustering performance against ground truth class labels.

### 4.1 Nucleus detection

Although we train the models on small human-annotatable images, we are eventually interested in inference on images of at least 100x larger size. In this case, the image resizer layer from the original Faster-RCNN architecture that resizes images to have the same shorter side makes nuclei in the training set look drastically larger in scale than those in inference set and hurts model performance. To tackle this, we replaced the original image resizer with a scaling layer that scales images by a constant factor irrespective of size.

We trained the models for a fixed number of steps (30000) using the hyper-parameters suggested in the Tensorflow Object Detection API. We increased maximum number of proposals during inference to accommodate larger images. Detection performance was measured by the mean average precision metric at threshold 0.50 (mAP@50).

### 4.2 Feature Extraction

ResNet-50 was used as first stage feature extractor of FasterRCNN. Features from block 1, block 2, and the convolutional ResNet-50 architecture were extracted and used in clustering. Note that in order to quantify clustering performance, we extracted features from ground truth bounding boxes in the validation set for which class labels are available. In real world applications, we expect the pipeline to be used for clustering nuclei detected by the model.

### 4.3 Nucleus clustering

We performed agglomerative clustering as implemented in scikit-learn [11] on the extracted features as well as the original RGB representations of nuclei. Clustering performance was measured using the cluster accuracy (purity) metric which measures the percentage of majority examples in the different clusters of a clustering [12]. This represents the classification accuracy one would achieve by labelling all nuclei in each cluster with the label of the majority class in that cluster.

## 5 EXPERIMENTS

Fine-tuning the pre-trained Faster-RCNN model on different datasets resulted in the following models:

- Model Zero: Faster RCNN model with Resnet-50 feature extractor pre-trained on COCO dataset.
- HER2: Model Zero fine-tuned on nucleus detection on Her2 data.
- DUALISH: Model Zero fine-tuned on nucleus detection on Dual ISH data.
- Model A: Model Zero fine-tuned on nucleus detection on Combined data.
- Model B: Model Zero fine-tuned on detection and classification on BB data.
- Model C: Model A fine-tuned on detection and classification on BB data.

### 5.1 Nucleus detection

A summary of detection (and classification, where class labels are present) performances of our models is given in Table 2. In Fig. 1, a few examples of detected vs. ground truth nuclei are presented. It can be seen that model A sometimes detects nuclei missed by the annotator and objects of ambiguous nature, resulting in model penalization.

**Table 2.** Nucleus detection and classification performance on held-out validation data.

| Validation Data | Model | mAP@50 |
| --- | --- | --- |
| Her 2 | HER2 | 78% |
| Dual ISH | DUALISH | 86% |
| Combined | Model A | 85% |
| BB | Model B | 70% |
| BB | Model C | 72% |

**Fig. 1.**
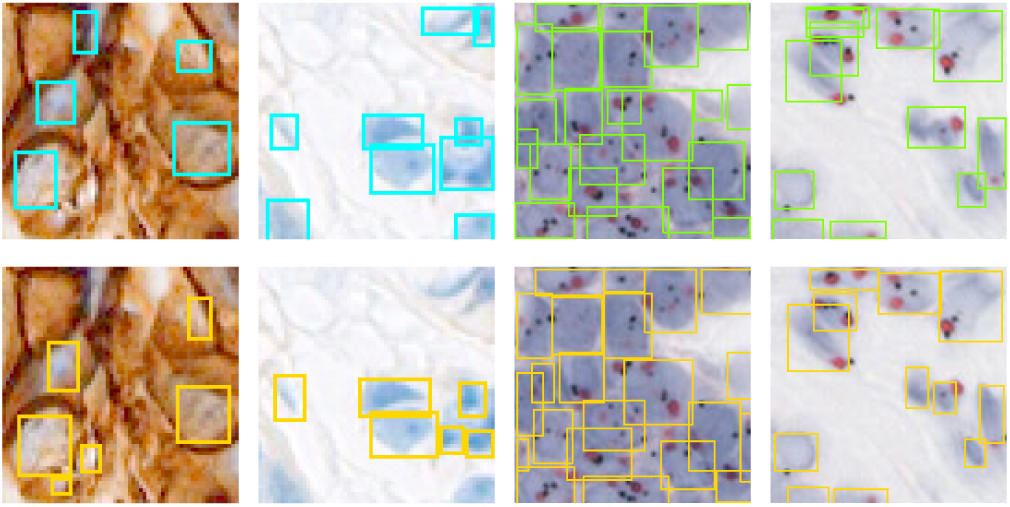
**Top:** Examples of nucleus detection results using model A on Her2 and Dual ISH images. **Bottom:** Ground truth annotations for the corresponding patches.

In Fig. 2, examples of nucleus detection results on large inference images are given. Such detected nuclei are used in later stages of the pipeline as described in the next subsections.

**Fig. 2.**
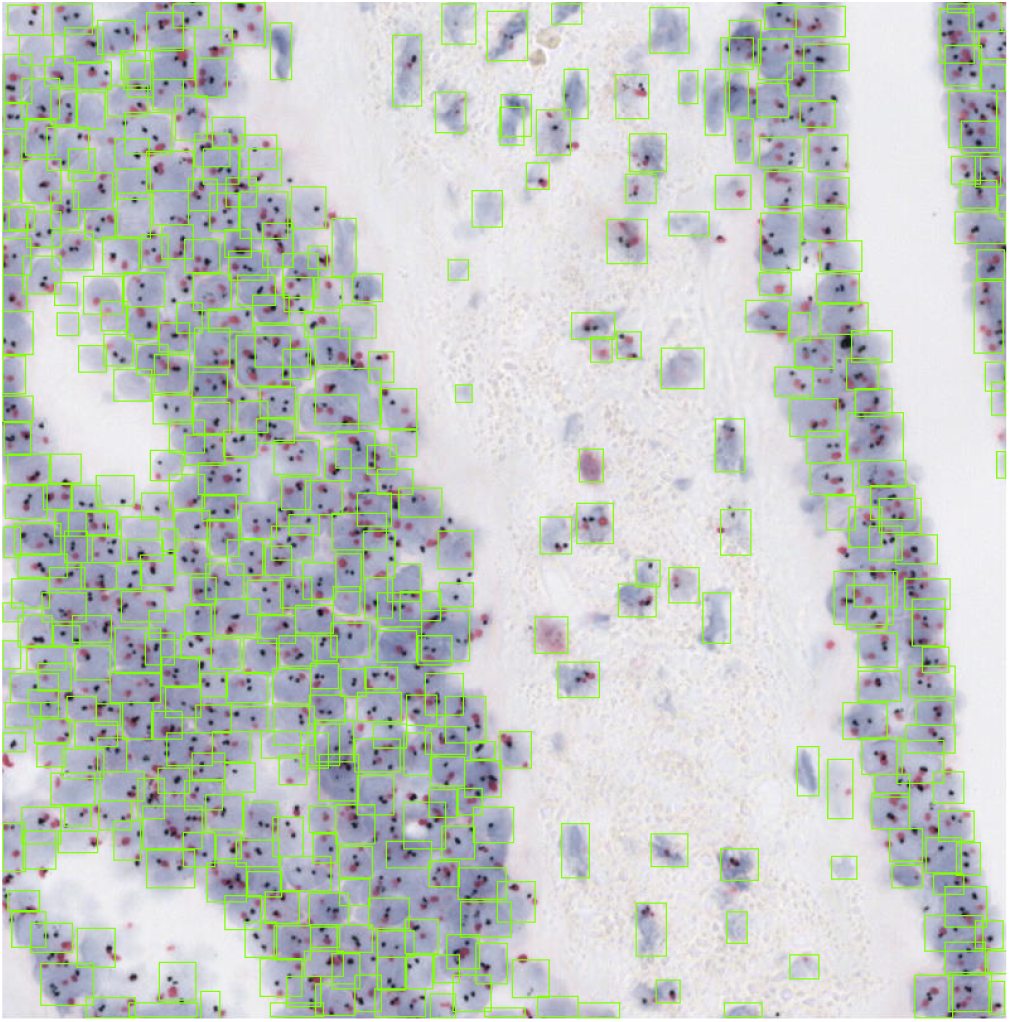
Model A inference on a 796×806 Dual ISH image.

### 5.2 Nucleus clustering

During the process of learning to detect and localize cells, Model A learns features that can facilitate clustering of nuclei. Although model A was not provided with any class labels, these features yield better clustering results compared to the original RGB values of cells (See Fig. 3).

**Fig. 3.**
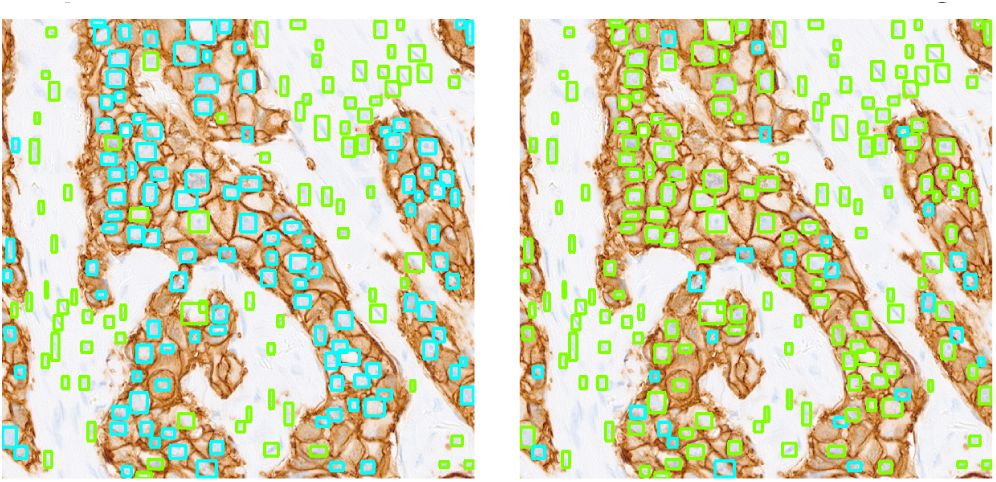
Qualitative nucleus clustering performance. **a.** Features extracted from block 1 of Model A enable meaningful clustering of nuclei. **b.** Clustering of cells based on their original RGB pixel values does not look meaningful.

### 5.3 Clustering of unseen nuclei classes

In this section, we investigate the transferability of class-agnostic nucleus detectors to clustering nucleus classes/stain types that the model has not seen during training. Fig. 4 summarizes clustering performance on BB data using representations extracted from different layers of Model A as well as the original RGB representation of nuclei. With only two clusters, Faster-RCNN features have a 8% advantage over RGB representation; This means if we cluster nuclei into 2 clusters using Faster-RCNN features and assign the majority class to all nuclei in each cluster, we get 98% classification accuracy compared to 90% if we use RGB representations instead.

**Fig. 4.**
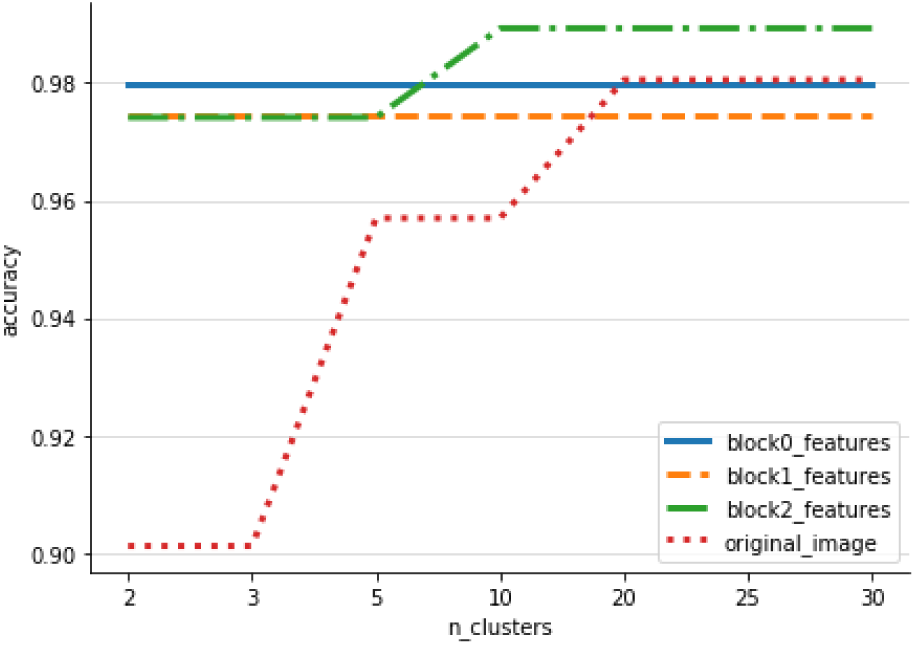
Evaluation of clustering based on Model A representation of BB validation data.

Does fine-tuning the model to detect and classify BB data increase the usefulness of the learned features for the task of clustering compared to the class-agnostic model? To answer this, we measured performance of clustering using features extracted from models Zero, A, B, and C. Although using representations learned by any of these models leads to at least 7% improvement over original RGB valules of nuclei, we observed less than 1% difference between results obtained by these models. This demonstrates that even the features learned from the COCO dataset can be used to meaningfully cluster nuclei and consequently ameliorate ground truth labelling even further.

### 5.4 Transferring further

BBYP images are stained by Discovery Yellow and Discovery Purple chromogens to identify 5 classes of cells of interest, i.e., ki67+/- tumor cells, Ki67+/- T-Cells and CD8+ T-Cells. Some of these classes are easily distinguishable by color, while others are defined by both color and morphology or context (See Fig. 5). We hypothesize that since CNN features provide a more global view of objects due to their multi-level abstraction compared to original image pixels, they should provide better distinction between nucleus classes that look similar in individual appearance but can be distinguished by contextual information. As shown in Figure 6, clustering BBYT nuclei using Model B’s features yields a 15% advantage over using original RGB representation of these nuclei, and continue to outperform them as we increase the number of clusters, supporting our initial hypothesis. Using features from other models led to similar results.

**Fig. 5.**
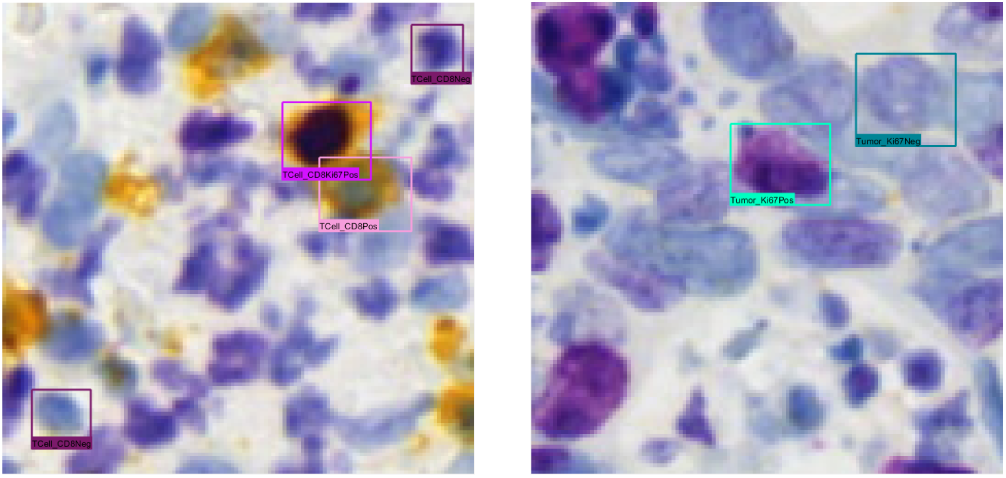
Examples of T-Cells (right) and tumor cells (left) in BBYP dataset. Ki67+ and Ki67-tumor cells have the same color as Ki67+ and Ki67-T-Cells, respectively. These cells can be differentiated based on cell size, shape, context, etc.

**Fig. 6.**
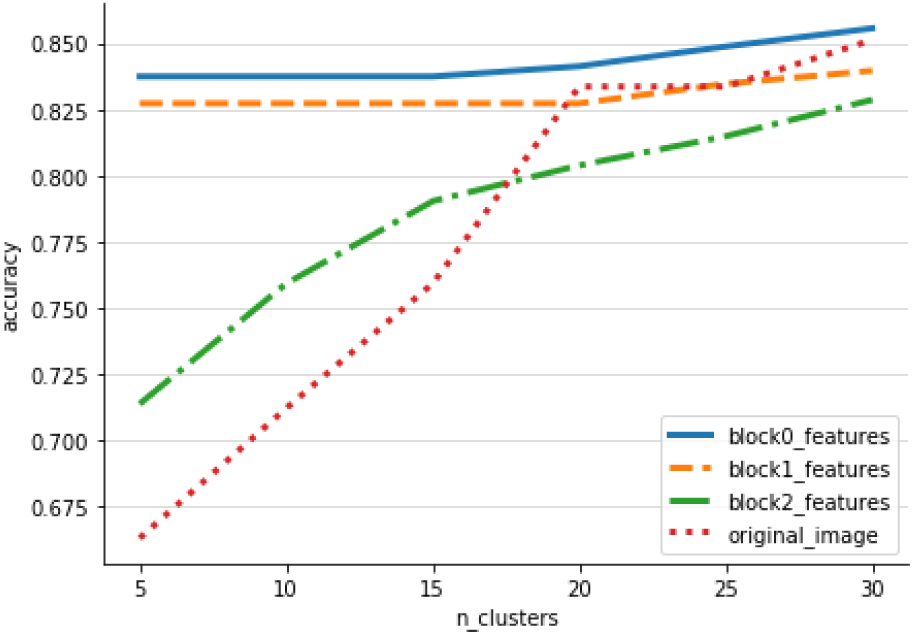
Evaluation of clustering based on representation of BBYP nuclei extracted from several layers of Model B.

## 6 CONCLUSION

Our proposed pipeline facilitates ground truth labeling by detecting nuclei in histopathology images and clustering detected nuclei into a few homogeneous subsets that can be annotated in bulk, reducing the burden of ground truth labeling from an order of number of cells in the dataset to the number of clusters.

We show that features learned from the COCO dataset are transferable for nucleus clustering and achieve over 98% cluster homogeneity on BB validation data. Using only 1800 bounding box annotations, we demonstrate that with the right training, Faster R-CNN can handle histopathology images of different magnifications and sizes at the same time, and perform inference on images 100x the size of training patches (the limitation here is not algorithmic but memory-related).

